# Development and Characterization of a *Sf-1*-Flp Mouse Model

**DOI:** 10.1101/2025.02.21.639566

**Authors:** Marco Galvan, Mina Fujitani, Samuel R. Heaselgrave, Shreya Thomas, Bandy Chen, Jenny J. Lee, Steven C. Wyler, Joel K. Elmquist, Teppei Fujikawa

**Affiliations:** Center for Hypothalamic Research, Department of Internal Medicine, UT Southwestern Medical Center, Dallas, Texas, USA; Department of Neuroscience, UT Southwestern Medical Center, Dallas, Texas, USA; Department of Pharmacology, UT Southwestern Medical Center, Dallas, Texas, USA; Peter O’Donnell Jr. Brain Institute, UT Southwestern Medical Center, Dallas, Texas, USA; Institute of Human Life and Ecology, Osaka Metropolitan University, Osaka, Japan

**Author notes:** Contribute equally (Marco Galvan, Mina Fujitani). Corresponding authors Joel K. Elmquist Center for Hypothalamic Research, Department of Internal Medicine, UT Southwestern Medical Center, 5323 Harry Hines Blvd, Dallas, Texas 75390-9077, Email), Teppei Fujikawa Address; Center for Hypothalamic Research, Department of Internal Medicine, UT Southwestern Medical Center, 5323 Harry Hines Blvd, Dallas, Texas 75390-9077, Email. **Disclosure:** The authors have nothing to disclose.

**Keywords:** SF-1, VMH, Flp-FRT

## Abstract

The use of genetically engineered tools, including combinations of Cre-LoxP and Flp-FRT systems, enable the interrogation of complex biology. Steroidogenic factor-1 (SF-1) is expressed in the ventromedial hypothalamic nucleus (VMH). Development of genetic tools, such as mice expressing Flp recombinase (Flp) in SF-1 neurons (*Sf-1*-Flp), will be useful for future studies that unravel the complex physiology regulated by the VMH. Here, we developed and characterized *Sf-1*-Flp mice and demonstrated its utility. Flp sequence was inserted into *Sf-1* locus with P2A. This insertion did not affect *Sf-1* mRNA expression levels and *Sf-1*-Flp mice do not have any visible phenotypes. They are fertile and metabolically comparable to wild-type littermate mice. Optogenetic stimulation using adeno-associated virus (AAV)-bearing Flp-dependent channelrhodopsin-2 (ChR2) increased blood glucose and skeletal muscle PGC-1α in *Sf-1*-Flp mice. This was similar to SF-1 neuronal activation using *Sf-1*-BAC-Cre and AAV-bearing Cre-dependent ChR2. Finally, we generated *Sf-1*-Flp mice that lack *β2-adrenergic receptors* (*Adrβ2*) only in skeletal muscle with a combination of Cre/LoxP technology (*Sf-1*-Flp::SKM^ΔAdrβ2^). Optogenetic stimulation of SF-1 neurons failed to increase skeletal muscle PGC-1α in *Sf-1*-Flp::SKM^ΔAdrβ2^ mice, suggesting that *Adrβ2* in skeletal muscle is *required* for augmented skeletal muscle PGC-1α by SF-1 neuronal activation. Our data demonstrate that *Sf-1*-Flp mice are useful for interrogating complex physiology.

## Introduction

The ventromedial hypothalamic nucleus (VMH) regulates multiple physiological responses and has been investigated for decades (1–3). For instance, the VMH regulates metabolism, food intake, behavior such as defensive behavior, reproduction, autonomic nervous system, and many other physiological functions (1–4). Although the role of the VMH in various physiological functions is well investigated, the precise molecular and neuronal mechanisms by which the VMH regulates physiology still remain largely unclear.

Steroidogenic factor-1 (SF-1, official gene name is *Nr5a1*) is expressed in the entire VMH early in embryonic development (5–7). In adults, SF-1 expression becomes more restricted to VMH neurons in the dorsomedial and center of the VMH (5, 6). Of note, SF-1 is also expressed in the pituitary gland, adrenal gland, and gonads like the testis and ovary (8, 9). We previously demonstrated that SF-1 neurons in the VMH (VMH^SF-1^ neurons) regulate skeletal muscle transcriptional events via the sympathoadrenal drive (10). Briefly, optogenetic stimulation of VMH^SF-1^ neurons using *Sf-1*-BAC-Cre (11) and adeno-associated virus (AAV) that contains Cre-dependent channelrhodopsin-2 (ChR2) (AAV-DIO-ChR2) (12) increases mRNA levels of skeletal muscle *peroxisome proliferator-activated receptor gamma coactivator 1-alpha* (PGC-1α, official gene name is *Ppargc1a*) (10). Surgical ablation of the adrenal gland or global knockout of *β2-adrenergic receptors* (*Adrβ2*) hampers inductions of skeletal muscle *Pgc-1α* expression, suggesting the sympathoadrenal drives and *Adrβ2* are required for VMH^SF-1^ neuronal regulation of skeletal muscle. Considering that *Adrβ2* is a major subtype of adrenergic receptors in skeletal muscle cells (13), VMH^SF-1^ neurons regulate skeletal muscle PGC-1α likely via direct actions on *Adrβ2* in skeletal muscle cells. However, *Adrβ2* is expressed in other metabolically important tissues, including the pancreas (14), liver (15), and VMH (16); thus it is still unclear which tissues are responsible for VMH^SF-1^ neuronal regulation of skeletal muscle *Pgc-1α* expression when using global *Adrβ2* KO mice. To determine whether *Adrβ2* in skeletal muscle cells is indeed *required* for VMH^SF-1^ neurons to regulate skeletal muscle PGC-1α, we needed to develop an animal model that facilitates studies to ablate *Adrβ2* only in skeletal muscle cells and simultaneously to manipulate VMH^SF-1^ neuronal activity. To this end, we developed mice expressing Flpe recombinase (Flp) in SF-1 neurons (*Sf-1*-Flp). Similar to the Cre-LoxP system, Flp recognizes FRT sites and induces a variety of recombinations depending on the directions and locations of FRT sites (17–20). Because there is minimum overlapping enzymatic recognition between Cre-LoxP and Flp-FRT systems (21), these systems can be used in combination, enabling complex genetic manipulations.

An insertion of Flp in *Sf-1* locus does not affect mRNA expression of *Sf-1* in the hypothalamus, pituitary gland, adrenal gland, and testis/ovary where SF-1 is expressed. We confirmed that *Sf-1*-Flp mice exhibit Flp activity in SF-1 neurons in the VMH without any visible phenotypes. Further, fundamental metabolism such as body weight and glucose levels in *Sf-1*-Flp mice is comparable to littermate wild-type control mice. By crossing *Sf-1*-Flp with mice lacking *Adrβ2* only in skeletal muscle by Cre-LoxP system, we achieve to activate VMH^SF-1^ neurons in mice lacking *Adrβ2* only in skeletal muscle. We found that lacking *Adrβ2* only in skeletal muscle cells hampers augmented skeletal muscle PGC-1α by VMH^SF-1^ neuronal activation. Collectively, our data demonstrate that *Sf-1*-Flp mouse is a valuable tool to decipher physiology that is regulated by VMH^SF-1^ neurons.

## Results

### Generation of *Sf-1*-Flp mice

We first attempted to insert Flpe recombinase, a temperature-stabilized form of Flp (22), into the downstream of *Sf-1* coding sequence region using either IRES or P2A. However, we could not obtain any founders with either method. SF-1 global knockout is lethal in mice (23). We concluded that inserting Flp into C terminal region of SF-1 disrupted *Sf-1* transcriptional activity by some unknown mechanisms, leading to the lethality. Thus, we decided to insert Flp upstream of the start codon of *Sf-1* with P2A, which is a self-cleaving peptide (24, 25) (**SFigure 1**). Using CRISPR/Cas9 approach as we described previously (26, 27), we successfully obtained *Sf-1*-Flp mice. We bred *Sf-1*-Flp mice with Ai65f^TB/TB^ mice, which bear Flp-dependent tdTomato in *Rosa26* locus (28) (*Sf-1*-Flp^+/-^::Ai65f^TB/-^). *Sf-1*-Flp^+/-^::Ai65f^TB/-^ mice expressed tdTomato in tissues where SF-1 is expressed such as the VMH, pituitary gland, adrenal gland, and gonads (**Figure 1A-F**). In the brain, tdTomato expression is restricted to the VMH and axons emanating from the VMH (**SFigure 2**). These expression patterns are consistent with previous reports (6, 29), suggesting no ectopic expression, at least in the brain. In line with previous studies (6, 11), a scatter of tdTomato-positive cells was observed outside of the VMH (**Figure 1A** and B, **SFigure2**). There are a couple of potential explanations for why these scatter tdTomato-positive cells were observed outside of the VMH. Although this may simply be an artifact due to the nature of transgenic animals, the following explanation is more likely the case. During embryonic development, SF-1 positive cells are migrated into the VMH from several areas such as median eminence (6). Some of these SF-1 cells during development fail to migrate into the VMH (6). Indeed, single-cell/nucleus RNA-sequence (sc/sn-RNAseq) studies have pointed out a small number of POMC-positive cells, which are located in the hypothalamic arcuate nucleus (ARH), expressing *Sf-1* (30). Further studies using cutting-edge technology, such as spatial RNA-sequence (31), will be warranted to determine genetic properties of these scatter cells outside of the VMH. Next, we determined whether the insertion of Flp disrupts expression of *Sf-1.* We found no differences in mRNA levels of *Sf-1* in the hypothalamus, pituitary gland, adrenal gland, and testis/ovary between heterozygous *Sf-1*-Flp mice and wild-type littermate control mice (**Figure 1G-K**). These data suggest that the insertion of Flp into 5’UTR of *Sf-1* locus unlikely affects *Sf-1* mRNA expression in *Sf-1*-Flp mice.

**Figure 1.**
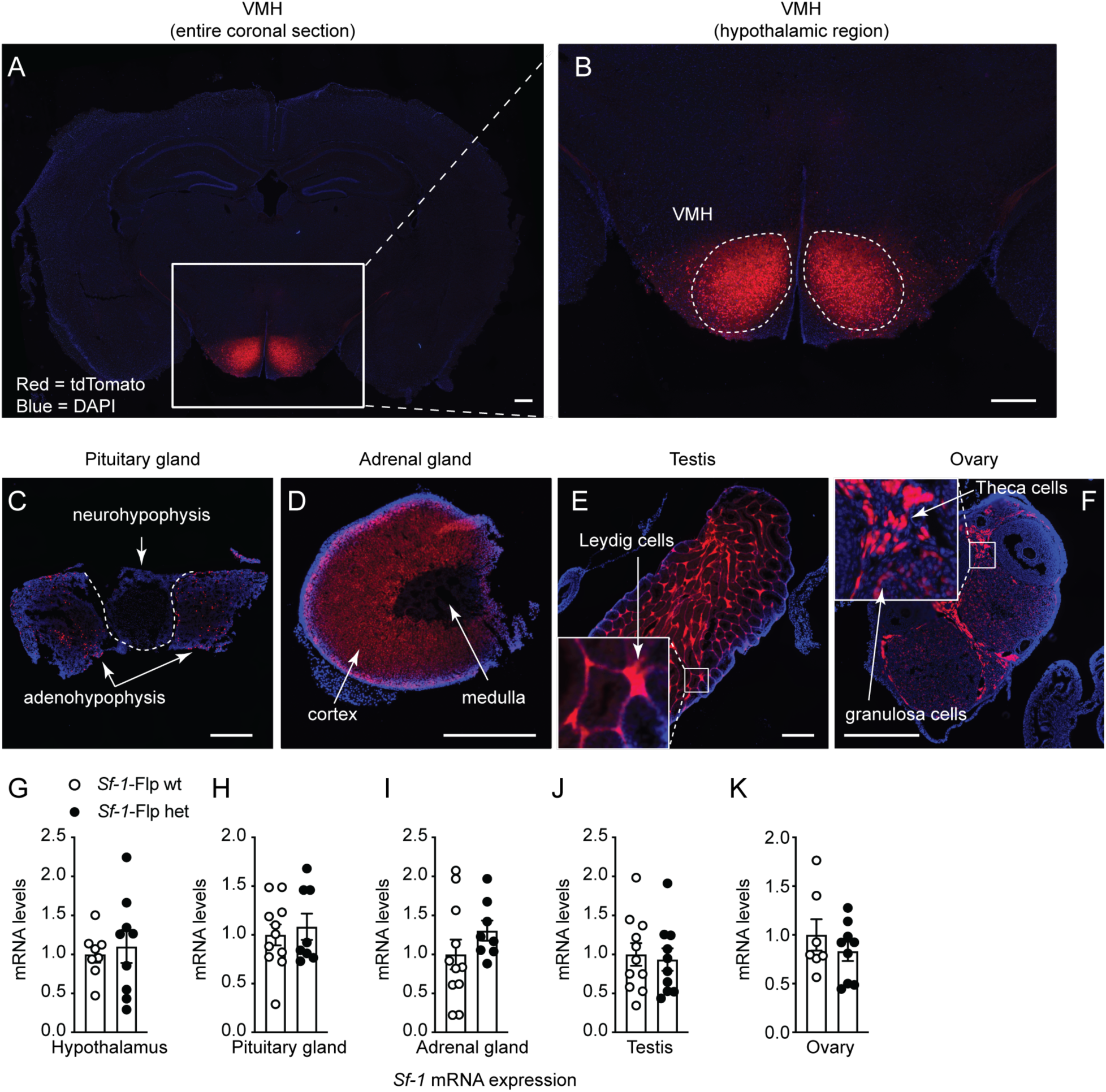
Generation of *Sf-1*-Flp mice. The representative figures of (**A and B**) the ventromedial hypothalamic nucleus (VMH), (**C**) pituitary gland, (**D**) adrenal gland, (**E**) the testis, and (**F**) ovary of *Sf-1*-Flp::Ai65f mice. Ai65f mice bearing Flp-dependent tdTomato in *Rosa26* locus. All white scale bars are 500 µm. mRNA levels of *Sf-1* in the (**G**) hypothalamus, (**H**) pituitary gland, (**I**) adrenal gland, (**J**) the testis in males, and (**K**) ovary in female of *Sf-1*-Flp heterozygous mice. The control group is composed of wild-type littermate mice. Values are mean ± S.E.M.

### *Sf-1*-Flp mice have normal body weight and whole body glucose metabolism

Next, we assessed baseline metabolic parameters in *Sf-1*-Flp mice by comparing heterozygous *Sf-1*-Flp mice with wild-type littermate control mice. Body weight, fat mass, and lean mass in *Sf-1*-Flp mice were comparable to those of control mice (**Figure 2A-H**). There were no significant differences in glucose tolerance and insulin tolerance between control and *Sf-1*-Flp mice (**Figure 2I, J, M, and N**). Further, fed and fast blood glucose levels were comparable in control and *Sf-1*-Flp mice (**Figure 2K, L, O, and P)**, suggesting that *Sf-1*-Flp mice do not have aberrant whole-body glucose metabolism. SF-1 is required for maintaining normal endocrine function in mice as *Sf-1* heterozygous global knockout mice exhibit aberrant blood leptin and corticosterone levels (32). Blood leptin, insulin, and corticosterone levels in fed and fasting status were comparable in control and *Sf-1*-Flp mice (**SFigure 3A-C**). We found that basal blood epinephrine levels in *Sf-1*-Flp male mice were significantly lower than in control mice (**Figure 3A**). In female mice, blood epinephrine levels tended to be lower than control mice, with the average value of epinephrine in *Sf-1*-Flp female mice being approximately 55% of the control group (**Figure 3C**). Blood norepinephrine levels were comparable between *Sf-1*-Flp and control mice (**Figure 3B and D**), suggesting that the synthesis of epinephrine was disrupted. Thus, we investigated whether epinephrine synthesis in the adrenal gland was disrupted by measuring mRNA levels of tyrosine hydroxylase (*Th*), dopamine-β-hydroxylase (*dβh*), and phenylethanolamine N-methyltransferase (*Pnmt*) in the adrenal gland, which are essential for epinephrine synthesis. mRNA levels of *Th*, *dβh*, and *Pnmt* in male adrenal glands tended to be lower than control mice (**Figure 3B-G**), and in females, they were significantly lower than control mice (**Figure 3H-J**). These results suggested that the synthetic pathway of epinephrine is disrupted by an insertion of P2A-Flp into the 5’-UTR of *Sf-1* locus. Overall, our data demonstrated that the insertion of P2A-Flp into the 5’-UTR of the *Sf-1* locus unlikely affects most SF-1 functions; however, it potentially has indirect effects on adrenal medulla functions.

**Figure 2.**
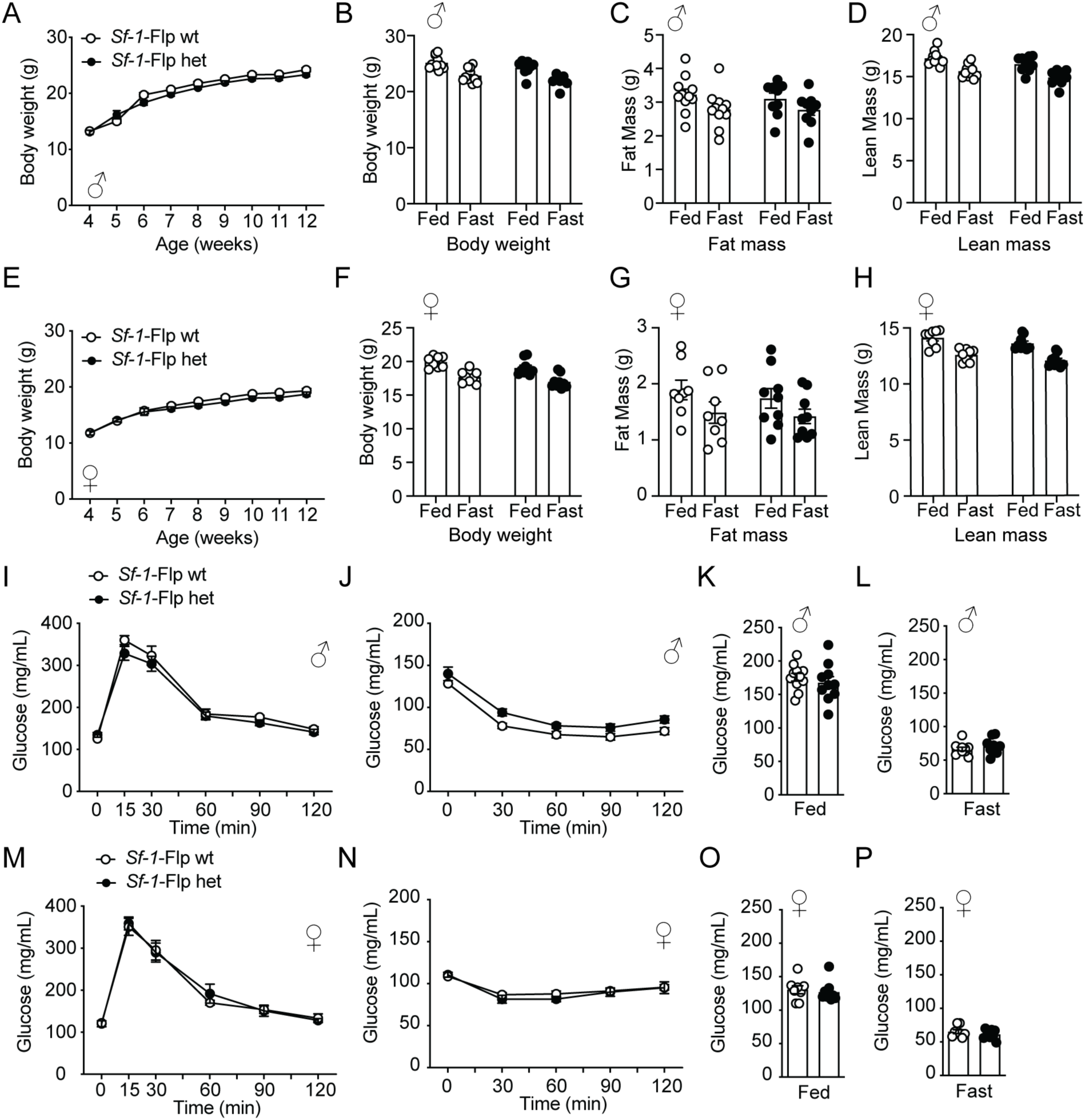
*Sf-1*-Flp mice do not have aberrant body weight and glucose metabolism. The time course of body weight of *Sf-1*-Flp heterozygous (**A**) male and (**E**) female mice. (**B, F**) Body weight, (**C, G**) fat mass, and (**D, H**) lean mass in fed and 24-hour fast conditions in *Sf-1*-Flp heterozygous (**B-D**) male and (**F-H**) female mice. Glucose tolerance test in *Sf-1*-Flp heterozygous (**I**) male and (**M**) female mice. Insulin tolerance test in *Sf-1*-Flp heterozygous (**J**) male and (**N**) female mice. Blood glucose levels in fed and 24-hour fast conditions in *Sf-1*-Flp heterozygous (**K and L**) male and (**O and P**) female mice. Values are mean ± S.E.M. Detailed statical analysis was described in Supplemetal Table 3.

**Figure 3.**
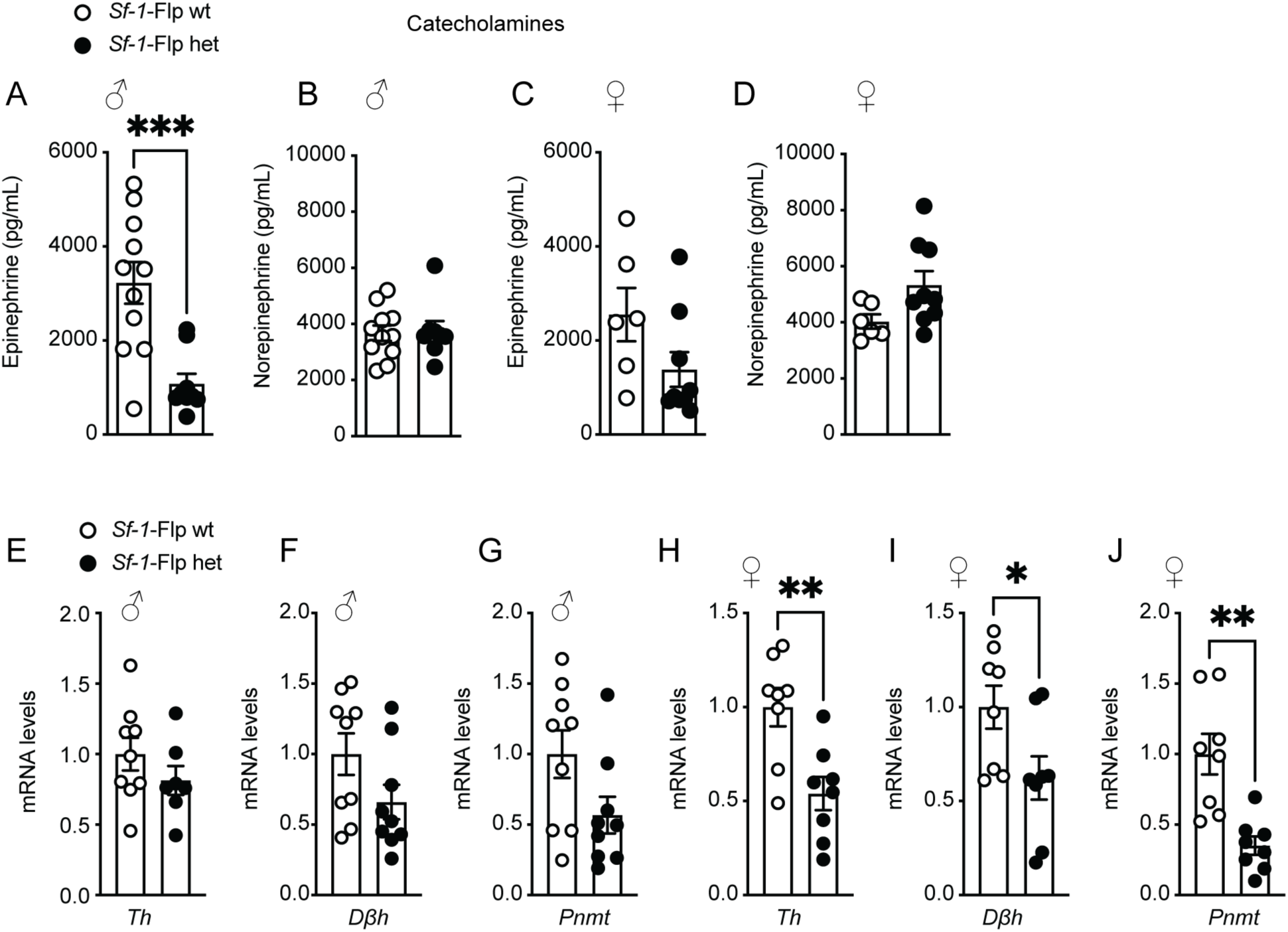
*Sf-1*-Flp mice have lower blood epinephrine. (**A-D**) Blood catecholamine levels in fed and 24-hour fast conditions in *Sf-1*-Flp heterozygous male and female mice. (**B-J**) mRNA levels of tyrosine hydroxylase (*Th*), dopamine-β-hydroxylase (*dβh*), and phenylethanolamine N-methyltransferase (*Pnmt*) in the adrenal gland in *Sf-1*-Flp heterozygous male and female mice. Values are mean ± S.E.M. *** p < 0.001, ** p <0.01, * p < 0.05.

### Optogenetic stimulation using *Sf-1*-Flp mice

Optogenetic stimulation of VMH^SF-1^ neurons using *Sf-1*-BAC-Cre mice and AAV-DIO-ChR2 increases blood glucose (10, 33) and mRNA levels of PGC-1α in skeletal muscle (10). Further, it suppresses appetite (34) and induces defensive behavior such as burst activities that include jumping and running (4, 35). To investigate whether we can utilize optogenetics in *Sf-1*-Flp mice, we delivered AAV bearing Flp-dependent ChR2 (AAV-fDIO-ChR2) into the VMH (*Sf-1*-Flp::ChR2) and examined whether optogenetic stimulation could produce a phenocopy of the stimulation in *Sf-1*-BAC-Cre mice. The control group received AAV that does not contain ChR2 but a fluorescent tag sequence (AAV-fDIO-EYFP) (*Sf-1*-Flp::EYFP). Optogenetic stimulation of VMH^SF-1^ neurons increased c-Fos in the VMH of *Sf-1*-Flp::ChR2 mice, suggesting that it is sufficient to induce augmented neuronal activities in VMH^SF-1^ neurons (**Figure 4C and D**). As expected, optogenetic VMH^SF-1^ neuronal activation in *Sf-1*-Flp::ChR2 mice increased blood glucose (**Figure 4E**), and burst activities were observed. We confirmed that the stimulation of VMH^SF-1^ neurons significantly increased blood epinephrine (**Figure 4F and G**) and mRNA levels of *Pgc-1α* in tibial anterior (TA) skeletal muscle of *Sf-1*-Flp::ChR2 mice (**Figure 4H**). We further examined whether optogenetic stimulation of VMH^SF-1^ neurons in *Sf-1*-Flp::ChR2 mice suppresses appetite. Mice were fasted overnight, which maximized their appetite. The stimulation significantly reduced food intake even though they were fasted overnight (**Figure 4I and J**). These results are a phenocopy of experiments using *Sf-1*-BAC-Cre and Cre-dependent AAV containing ChR2 (10). Overall, these data indicate that optogenetic tools can be used in *Sf-1*-Flp mice.

**Figure 4.**
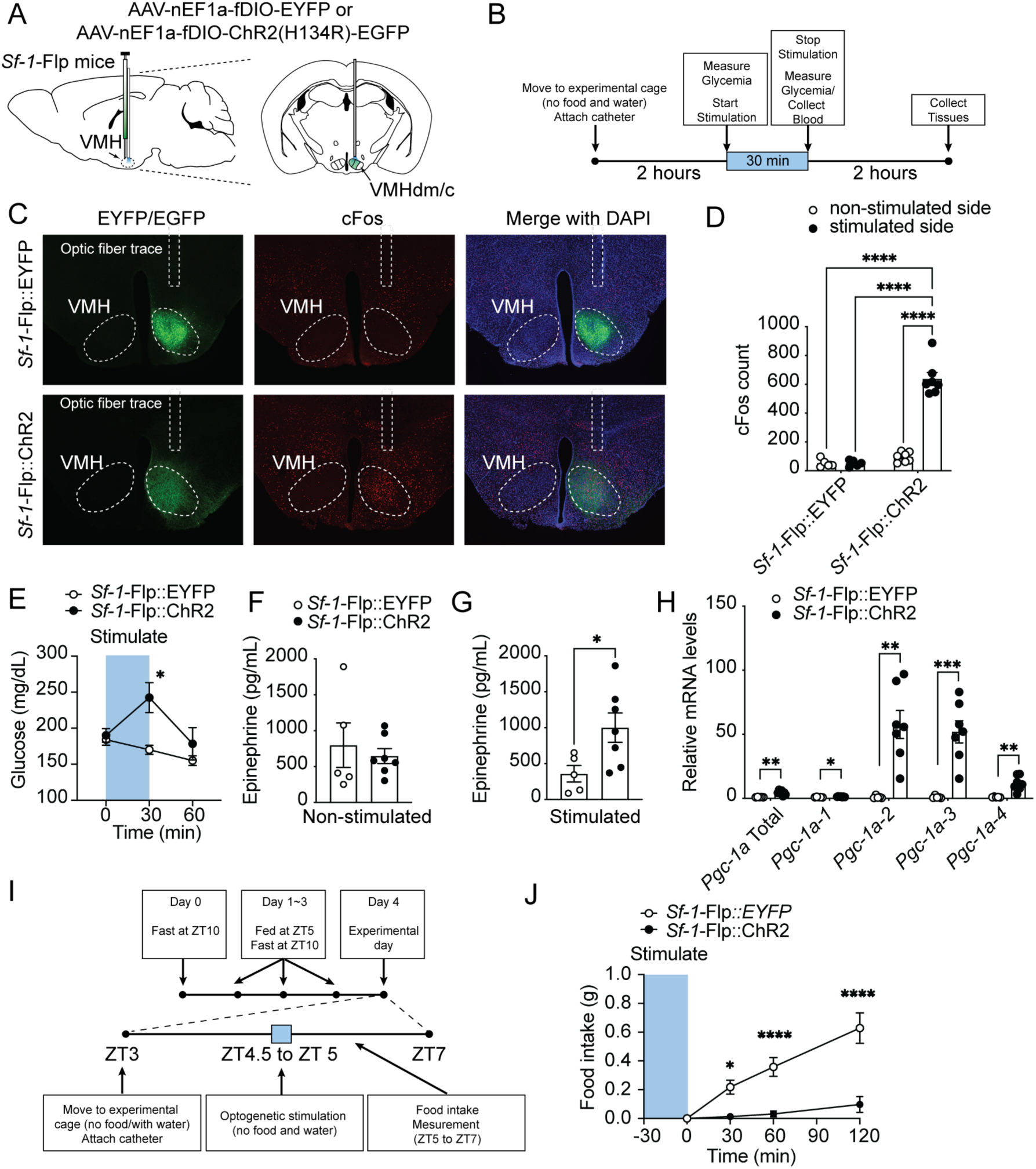
Optogenetic activation of VMH^SF-1^ neurons using *Sf-1*-Flp mice. (**A**) Schematic figure of targeting site of VMH^SF-1^ neurons using *Sf-1*-Flp mice and AAV containing Flp-dependent ChR2. (**B**) Experimental design and stimulus setting of optogenetics. (**C**) Representative figures of c-Fos expression pattern in the hypothalamus of mice expressing EGFP (*Sf-1*-Flp::EYFP) or ChR2 (*Sf-1*-Flp::ChR2) in VMH^SF-1^ neurons after optogenetic stimulation. (**D**) Number of c-Fos positive cells in the VMH of *Sf-1*-Flp::ChR2 mice after optogenetic stimulation. (**E**) Blood glucose levels in *Sf-1*-Flp::ChR2 mice with optogenetic stimulation. (**F and G**) Blood catecholamine levels and (**H**) mRNA expression levels of *Pgc1*-α isoform in skeletal muscle of *Sf-1*-Flp::ChR2 mice after optogenetic stimulation. (**I**) Experimental design for food intake study. (**J**) Food intake in *Sf-1*-Flp::ChR2 mice with optogenetic stimulation. All mice were male. Values are mean ± S.E.M. *** p < 0.001, ** p <0.01, * p < 0.05.

### Combination of Cre-LoxP and Flp-FRT system in *Sf-1*-Flp mice

Finally, we tested whether *Adrβ2* in skeletal muscle cells is *required* for the induction of skeletal muscle *Pgc-1α* mRNA by VMH^SF-1^ neuronal activation. To do so, we crossed *Sf-1*-Flp to mice expressing tamoxifen-inducible Cre under the control of the human skeletal muscle cell promoter (HAS-MCM) (36) and *Adrβ2* floxed (37) mice (*Sf-1*-Flp::SKM^ΔAdrβ2^). As previous studies showed (38), we confirmed that Cre activity was induced in skeletal muscle of HAC-MCM mice by tamoxifen-containing (TAM) diet (**SFigure 4A**). Control group was composed of littermate *Sf-1*-Flp^+/-^::*Adrβ2*^flox/flox^ mice (control). All mice were fed with TAM diet. After recovery from TAM diet, AAV-fDIO-EYFP or AAV-fDIO-ChR2 was microinjected into the VMH of control or *Sf-1*-Flp::SKM^ΔAdrβ2^ mice (Control::EYFP, Control::ChR2, *Sf-1*-Flp::SKM^ΔAdrβ2^::EYFP, and *Sf-1*-Flp::SKM^ΔAdrβ2^::ChR2) (**Figure 5A**). Deletion of *Adrβ2* in skeletal muscle cells did not affect increased blood glucose levels by VMH^SF-1^ neuronal activation (**Figure 5B**). In line with **Figure 4**, optogenetic VMH^SF-1^ neuronal activation in Control::ChR2 mice significantly increased skeletal muscle *Pgc-1α* mRNA than Control::EYFP (**Figure 5C-H**), whereas deletion of *Adrβ2* in skeletal muscle cells significantly hampers augmented skeletal muscle *Pgc-1α* mRNA by VMH^SF-1^ neuronal activation (**Figure 5C-H**). Of note, TAM diet significantly reduced mRNA levels of *Adrβ2* in skeletal muscle of *Sf-1*-Flp::SKM^ΔAdrβ2^, but not that of control mice (**SFigure 4B**). These data demonstrate that *Adrβ2* in skeletal muscle cells is *required* for augmented skeletal muscle *Pgc-1α* mRNA by VMH^SF-1^ neuronal activation. In addition, these results articulate that *Sf-1*-Flp mice can be used with combinations of Cre-LoxP system.

**Figure 5.**
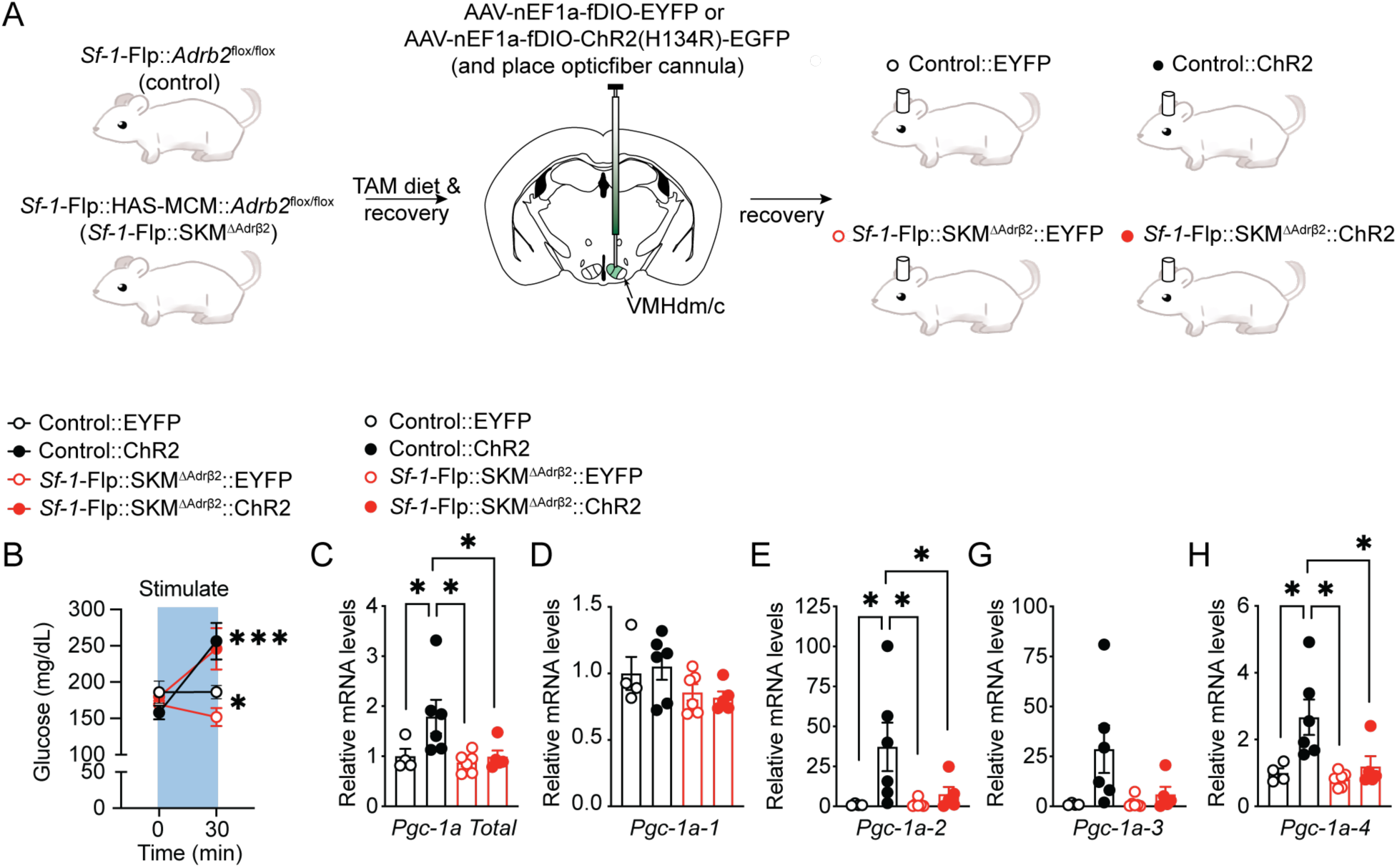
*Adrβ2* in skeletal muscle cells is *required* for augmented skeletal muscle PGC-1α mRNA by VMH^SF-1^ neuronal activation. (**A**) Experimental design to generate mice lacking *Adrb2* in skeletal muscle cells (*Sf-1*-Flp::SKM^ΔAdrβ2^) and expressing ChR2 in VMH^SF-1^ neurons (*Sf-1*-Flp::SKM^ΔAdrβ2^::ChR2). AAV-fDIO-EYFP was used as control-AAV for AAV-fDIO-ChR2 (*Sf-1*-Flp::SKM^ΔAdrβ2^::EYFP). Control group that do not lack *Adrb2* in skeletal muscle cells (control) was also administered either AAV-fDIO-EYFP or AAV-fDIO-ChR2 (Control::EYFP or Control::ChR2). (**B**) Blood glucose levels of *Sf-1*-Flp::SKM^ΔAdrβ2^::ChR2 mice after optgenetic stimulation. (**C-H**) mRNA expression levels of *Pgc1*-α isoform in skeletal muscle of *Sf-1*-Flp::SKM^ΔAdrβ2^::ChR2 mice after optogenetic stimulation. All mice were male. Values are mean ± S.E.M. *** p < 0.001, * p < 0.05.

## Discussion

In this study, we generate *Sf-1*-Flp mice that express Flpe recombinase in SF-1 cells and demonstrate that *Sf-1*-Flp mice can be useful in investigating complex physiology. Our previous report indicates that *Adrβ2* contributes to VMH^SF-1^ neuronal regulation of skeletal muscle transcriptional events (10). However, it was unclear whether VMH^SF-1^ neurons regulate a direct action of *Adrβ2* in skeletal muscle cells because *Adrβ2* is widely expressed throughout the body in both rodents and humans (39–41). Our current study demonstrates that *Adrβ2* in skeletal muscle cells is indeed *required* for augmented skeletal muscle *Pgc-1α* mRNA by VMH^SF-1^ neuronal activation. This strongly suggests that a direct action of *Adrβ2* in skeletal muscle cells is key for VMH^SF-1^ neuronal regulation of skeletal muscle transcriptional events and functions.

VMH^SF-1^ neurons contribute to beneficial effects of exercise including reductions in fat mass (42). Previous studies indicate that VMH^SF-1^ neurons are key to exercise-induced skeletal muscle adaptations via the sympathetic nervous system (10, 42). Our data (**Figure 5**) pinpoint that *Adrβ2* in skeletal muscle cells is likely required for VMH^SF-1^ neurons to regulate skeletal muscle function, although further studies are needed to elucidate what precise physiological functions VMH^SF-1^ neurons contribute to regulate in the context of long-term exercise training. These functions may include the mitochondrial respiratory system and switching fiber-type composition. For instance, long-term exercise training has been shown to improve mitochondrial function and alter muscle fiber-type (43–45). If VMH^SF-1^ neuronal activation is repeated for a certain duration (e.g., 8 to 12 weeks), it may enhance mitochondrial function and influence fiber-type components as recurrent exercise training does.

There are a couple of technical notes to be aware of for future studies when *Sf-1*-Flp mice are used. First, although we examined Flp activity in *Sf-1*-Flp mice throughout the brain (**SFigure 2**), we did not survey all peripheral tissues. As the nature of genetically-engineered mice, it is still possible that *Sf-1*-Flp mice may have ectopic Flp expression in some tissues where endogenous SF-1 is not expressed. Further, we used tdTomato expression as a surrogate of Flp activity, but it does not guarantee that Flp works with other genetically-engineered tools such as transgenic mice or AAV that bear FRT sites. Thus, it is important to validate each experimental model. Second, we found that *Sf-1*-Flp mice have lower blood epinephrine levels (**Figure 3**). Because *Sf-1* mRNA levels in the adrenal gland are comparable between *Sf-1*-Flp and control mice (**Figure 1**) and *Sf-1* is only expressed in the cortex of the adrenal gland, not the medulla where catecholamines are produced, it is unclear how the insertion of P2A-Flp into 5’-UTR disrupts the production of epinephrine in the adrenal gland. Optogenetic studies show that *Sf-1*-Flp mice can increase blood epinephrine levels by VMH^SF-1^ neuronal activation, suggesting that *Sf-1*-Flp mice can modify epinephrine releases upon afferent neuronal inputs. Nonetheless, we suggest that it is ideal for studies to contain a control *Sf-1*-Flp group for proper interpretation of the results. Further investigations will be warranted on the mechanism underlying the effects of the transgenic insertion into 5’UTR of *Sf-1* locus on blood epinephrine levels in *Sf-1*-Flp mice. Third, we noticed when AAV-fDIO-ChR2 was administered into the VMH of *Sf-1*-Flp mice, it took at least 5 to 6 weeks to have metabolic phenotype induced by VMH^SF-1^ neuronal activation (e.g., increased glucose levels). When we use *Sf-1*-BAC-Cre mice and AAV-DIO-ChR2 (10), it only takes 2 to 3 weeks to observe a metabolic phenotype. The reason for these discrepancies of durations is unclear. However, it is important to keep this in mind when AAV approaches are used in *Sf-1*-Flp mice.

*Sf-1*-Flp mice will provide unique opportunities for those who investigate the physiology of SF-1 cells, especially SF-1 neurons in the VMH. As shown in **Figure 5**, *Sf-1*-Flp mice unlock the possibilities of conducting experiments that manipulate genes in peripheral cells and modulate SF-1 neuronal activity or their function simultaneously. Further, *Sf-1*-Flp mice can be used for temporal conditioned manipulation combined with AAV approach such as using AAV bearing Flp-dependent Cre recombinase (AAV-fDIO-Cre). For instance, leptin receptors (LEPR) in the VMH appear to be important in regulating metabolism (11, 46), but it may be intriguing to determine the role of LEPR in the VMH in adults. If AAV-fDIO-Cre is administered into the VMH of *Sf-1*-Flp mice bearing two LoxP sites in LEPR locus (47) (*Sf-1*-Flp::*Lepr*^flox/flox^ mice), we may be able to achieve knockdown of LEPR specifically in VMH^SF-1^ neuron in adults. These approaches enable circumventing potential developmental compensations that happen when conventional Cre lines are used (48) and may unravel the role of interested genes in SF-1 neurons in adults.

In addition, *Sf-1*-Flp mice will enable the investigation of the detailed function of subpopulation in SF-1 neurons. Studies using sc/sn-RNAseq reveal genetically distinct subpopulations of the VMH, suggesting that the VMH is an extremely heterogeneous nucleus (16, 49). To interrogate which neuronal subpopulations of the VMH contribute to different physiological functions, *Sf-1*-Flp mice can be valuable as it can be applied such as using the INTERSECT method (21). For instance, LEPR is widely expressed in the hypothalamus, including the ARH or dorsomedial hypothalamic nucleus (DMH) (50, 51).

The ARH and DMH are juxtaposed to the VMH, thus it is technically difficult to target LEPR only in the VMH by microinjections or other injection methods. Using INTERSECT method in mice expressing Flp in SF-1 neurons and Cre in LEPR-expressing neurons by crossing *Sf-1*-Flp and *Lepr*-IRES-Cre mice (52), targeting LEPR neurons only in the VMH can be achieved.

In conclusion, our data demonstrate that *Sf-1*-Flp mice can be a great asset for neuroscience, metabolism, and many other fields that investigate the roles of cells expressing SF-1.

## Methods

### Sex as a biological variable

Both male and female mice were used in Figure 1-3. Male mice were used in optogenetic studies (Figure 4 and Figure 5).

### Generation of *Sf-1*-Flp mice

We utilized CRISPR/Cas9 approach to generate *Sf-1*-Flp mice. The sequence of *Mus musculus Nr5a1* (ENSMUST00000028084.5) was obtained from the Ensembl Genome Database (53, 54). Exon 2 of the *Nr5a1* gene encoding the translational start sight was targeted using the following guide RNA: 5’-GUACGAAUAGUCCAUGCCCGGUUUUAGAGCUAUGCU-3’; For the HDR repair template a 1499 bp IDT single-stranded DNA Megamer^TM^ (Intergrated DNA Technologies, US) was used encoding the Flpe recombinase sequence immediately followed by the P2A self-cleaving peptide sequence (22, 25). 5’ and 3’ homology arms were 45 and 92 bp respectively. Guide RNA, trcRNA, Cas9 protein, and HDR Template (all from IDT) were administered through a pronuclear injection in C57Bl/6NCrl (Charles River, US) zygotes by the UT Southwestern Transgenic Technology Center. A schematic design of *Sf-1*-Flp insertion was described in **SFigure 1**. Founders were screened by PCR followed by Sanger sequencing. To validate Flp activity *in situ*, we bred founders with Ai65f mice that express Flp-dependent tdTomato in *Rosa26* locus (28). Importantly, *Sf-1*-Flp mice will be publicly available upon the materials transfer agreement between the University of Texas Southwestern Medical Center (UTSW) and the recipient institute.

### Genetically-engineered mice and mouse husbandry

*Sf-1*-Flp mice were generated as outlined above. HAS-MCM (RRID:IMSR_JAX:025750) (36), Ai65f (RRID:IMSR_JAX:032864) (28), and Ai14 (RRID:IMSR_JAX:007914) (55) mice were purchased from the Jackson Laboratory. *Abrb2* floxed mice (15) were obtained from Dr. Florent Elefteriou at Baylor College of Medicine with the permission of Dr. Gerard Karsenty at Columbia University. Ear or tail gDNA was collected from each mouse to determine its genotype. A KAPA mouse genotype kit was used for PCR genotyping (Roche, US). The sequences of genotyping primers and expected band sizes for each allele are described in **Supplemental Table 1**. Mice were housed at room temperature (22–24 C°) with a 12 hr light/dark cycle (lights on at 6am, and 7am during daylight saving time) and fed a normal chow diet (2016 Teklad global 16% protein rodent diets, Envigo, US). Magnetic-resonance whole-body composition analyzer (EchoMRI, TX US) was used to analyze the fat and lean mass of mice. Mice were maintained in groups and singly housed after adeno-associated virus (AAV) injections and optic fiber probes insertions. Animal care was according to established NIH guidelines.

### AAV injections and optic fiber probe insertions

rAAV5-nEF1α-fDIO-hChR2(H134R)-EGFP-WPRE-hGH polyA, (Catalog# PT-1384; 5.5 x 10^12^ VM/mL) and rAAV-nEF1α-fDIO-EYFP-WPRE-hGH polyA (Catalog# BHV12400246; 2.8 x 10^12^ VM/mL) were purchased from Biohippo (MD, USA). AAVs were unilaterally administered into the right side of the VMH of mice using a UMP3 UltraMicroPump (WPI, US) with 10 µL NanoFil microsyringe (WPI) and NanoFil needle (WPI; Catalog# NF33BV). The volume of AAVs was 500 nL at the rate 100 nL per minute, and the needle was left for another 5 minutes after the injection was finished. The coordinates of VMH-microinjection were AP; −1.4 L; +0.5, and D; −5.5 (from Bregma). The optic fiber probe was inserted as following coordinates; AP; −1.4, L; +0.5, and D; −5.0. The configuration of the probe was 200 µm Core, 0.39NA, Ø2.5 mm ceramic ferrule, and 6mm length (RWD Life science Inc, US). The fiber probe was secured by adhesion bond (Loctite 454, Loctite Inc, US). Mice were allowed to recover for at least five to six weeks after the AAV injections to express recombinant proteins fully.

### Optogenetics

As we previously reported (10), the laser unit (Opto engine LLC, US; Catalog# MDL-III-470) was used for VMH^SF-1^ neuronal stimulations. The power of tips was set to ∼1 mW/mm^2^. The customed transistor-transistor logic generator was built based on the design by the University of Colorado Optogenetics and Neural Engineering Core, which is supported by the National Institute of Neurological Disorders and Stroke of the National Institutes of Health under award number P30NS048154. Rotary joint patch cable (Thorlabs; Catalog# RJPSF4 for LED and RJPFF4 for the laser) was used to connect to the laser unit. The quick-release interconnect (Thorlabs; Catalog# ADAF2) was used to connect the rotary joint patch cable to the fiber probe attached to the mouse head. The stimulation was set to; 5 ms duration, 20 Hz, 2 seconds activation and 2 seconds rest cycle, 30 minutes.

### Assessment of food intake

Mice were singly housed for food intake measurement. As shown in Figure 3H, mice are acclimated to time-restricted food availability for 3 days. Water was available throughout acclimation and experimental days except for when 30 minutes of optogenetic stimulation was executed. Food was available between ZT5 to ZT10. On the experimental day, mice were moved to the experimental cages without food. After optogenetic stimulation was finished at ZT5, food and water were provided. Food weight was measured at 30, 60, and 120 minutes.

### Immunohistochemistry and Fos counting

Mouse brains were prepared as previously described (10, 56). Anti-c-Fos (Abcam, US; Catalog# ab190289, Lot# GR3418522-1, and RRID:AB_2737414) and secondary fluorescent antibodies (Thermo Fisher Scientific, Inc, US; Catalog# A-21203, Lot#WD319534, and RRID:AB_141633) were used. Dilution factors for antibodies were 1:2000 for the primary and 1:200 for the secondary antibodies. Images were captured by a fluorescence microscopy (Leica Inc, US; Model DM6 B) or a slide scanner (Axioscan 7; (Zeiss US, US). Quantification of c-Fos-positive cells was described previously (10). Briefly, the exposure of captured images was adjusted and the VMH region was clipped by Photoshop based on the mouse brain atlas [28]. Clipped images were exported to Fiji, and the number of cells expressing Fos was counted by particle measurement function.

### Assessment of glucose, catecholamines, and hormone levels in the blood

Blood glucose was measured by a glucose meter as previously described (42, 57, 58). Blood for leptin, insulin, and corticosterone measurement was collected by tail-nicking with a heparin-coated capillary glass tube. We removed food 2 hours before blood collection to avoid immediate postprandial effects on blood metabolite and hormone and defined this condition as fed status in this study. For catecholamine measurement, trunk blood (**Figure 3A**) or submandibular blood (**Figure 4F**) was used. Plasma catecholamines and hormone levels were measured as previously described (42, 58). Briefly, the plasma catecholamines were analyzed by the Vanderbilt Hormone Assay & Analytical Services Core. Plasma leptin (Crystal Chem Inc, US; Catalog# 90030 and RRID:AB_2722664), insulin (Crystal Chem Inc, US; Catalog# 90080 and RRID:AB_2783626), and corticosterone (Cayman Chemical, US; Catalog #501320 and RRID:AB_2868564) levels were determined by commercially available ELISA kits.

### Glucose and Insulin tolerance tests

For glucose tolerance test, mice were fasted for 6 hours prior to testing starting at ZT2 with water provided ad libitum. For the glucose tolerance test, mice received an intraperitoneal injection of glucose solution (2 g/kg per BW). Glucose solution was purchased from ThermoFisher (catalog # A2494001). For insulin tolerance test, mice were fasted for 4 hours prior to testing starting at ZT4 with water provided ad libitum. Mice received i.p. insulin (Humalin® R, Eli Lilly, US) at 0.75 U/kg per BW for male mice and 0.50 U/kg per BW for female mice. Blood glucose was measured using a handheld glucometer (Bayer’s Contour Blood Glucose Monitoring System; Leverkusen Germany).

### Assessment of mRNA

mRNA levels in tissues were determined as previously described (42, 59). The sequences of primers for qPCR using SYBR or the catalog of Taqman probes are described previously (60) and in **Supplemental Table 2**.

### Graphic software, Data analysis, and statistical design

The schematic design of *Sf-1*-Flp mice in **Supplemental Figure 1** was generated by SnapGene (GSL Biotech LLC, US). The data are represented as means ± S.E.M. GraphPad PRISM version 10 (GraphPad, US) was used for the statistical analyses and *P*<0.05 was considered as a statistically significant difference. A detailed analysis of each figure was described in **Supplemental Table 3**. The sample size was decided based on previous publications (26, 42, 56–59, 61–63), and no power analysis was used. Figures were generated using PRISM version 10, Illustrator 2023 and 2025, and Photoshop 2023 and 2025 (Adobe Inc, US).

### Study approval

All procedures using mice were approved by the Institutional Animal Care and Use Committee of UTSW, and animal experiments were performed at UTSW.

### Data availability

All datasets generated during the current study are provided in **the Supporting Data Values File**.

## Acknowledgment

We would like to thank Dr. Florent Elefteriou at Baylor College of Medicine and Dr. Gerard Karsenty at Columbia University for sharing *Abrb2* floxed mice. We would like to thank UTSW Whole Brain Microscopy Facility (RRID:SCR_017949). Equipment of the UTSW Whole Brain Microscopy Facility used in this research was purchased with the NIH award (1S10OD032267-01 to Dr. Denise Ramirez). This study was supported by the National Institute of Health (To J.K.E., P01DK119130, R01DK088423, and R01DK100659, UTSW NORC P30DK127984), and by JSPS KAKENHI Grant Number 24K23124 to T.F.

## Contribution

M.G. and M.F. designed, performed, and analyzed experiments, and edited the manuscript. M.G. and M.F. equally contributed to this study, and they have the right to list them as the first author. S.R.H., S.T., B.C., and J.J.L. performed experiments. S.C.W designed *Sf-1*-Flp materials such as gRNA and edited the manuscript. J.K.E. supervised and edited the manuscript. T.F. designed, performed, supervised, and analyzed experiments, and wrote and finalized the manuscript.

## Supplemental Figure Legends

**Supplemental Figure 1.**
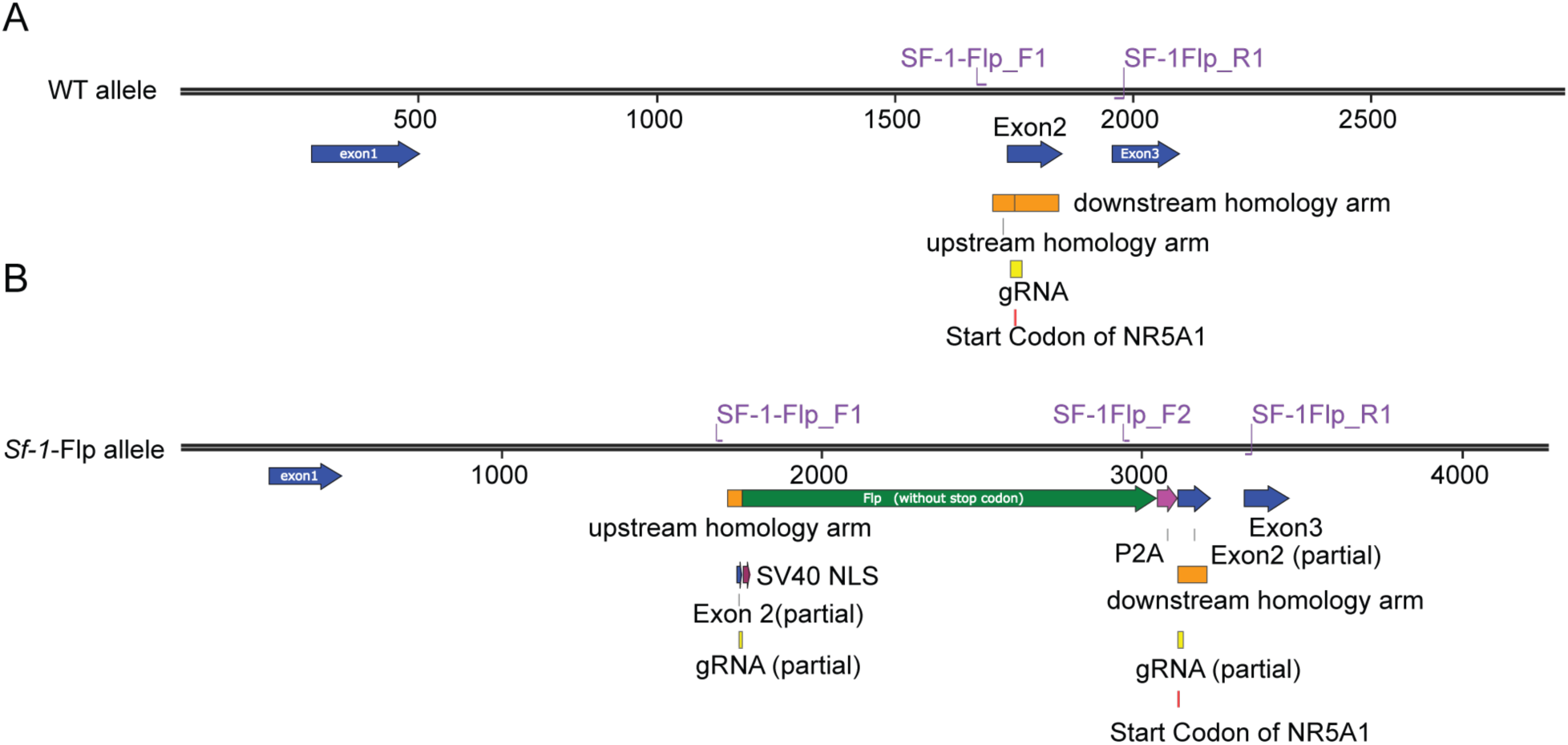
The schematic design of *Sf-1*-Flp mice. WT allele (upper) and *Sf-1*-Flp allele (lower) sequences are described. SF-1-Flp_F1, F2, and R3 indicate the location of genotype primers for *Sf-1*-Flp mice.

**Supplemental Figure 2.**
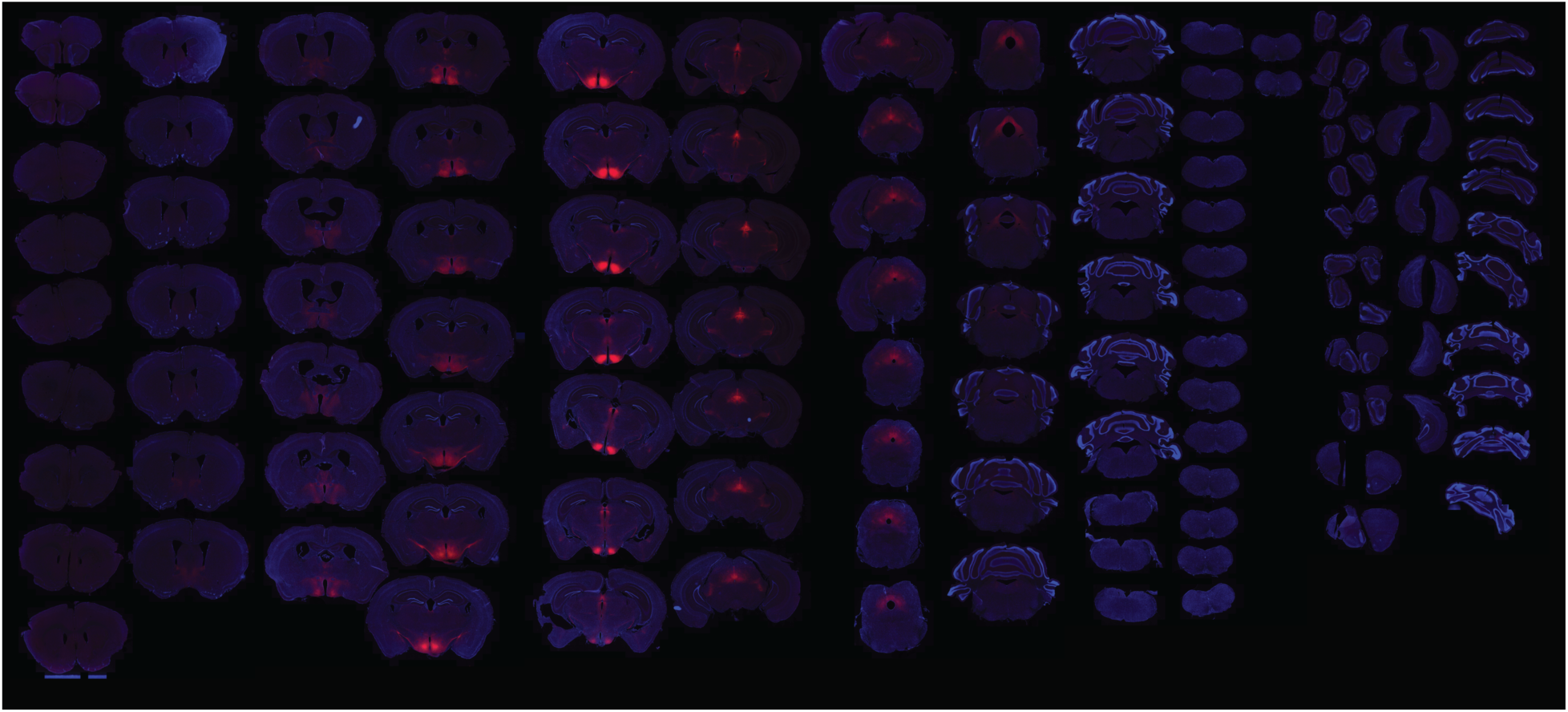
Brains sections of *Sf-1*-Flp::Ai65f^TB/-^ mice. Blue color indicates DAPI staining and red color indicates SF-1 positive cells.

**Supplemental Figure 3.**
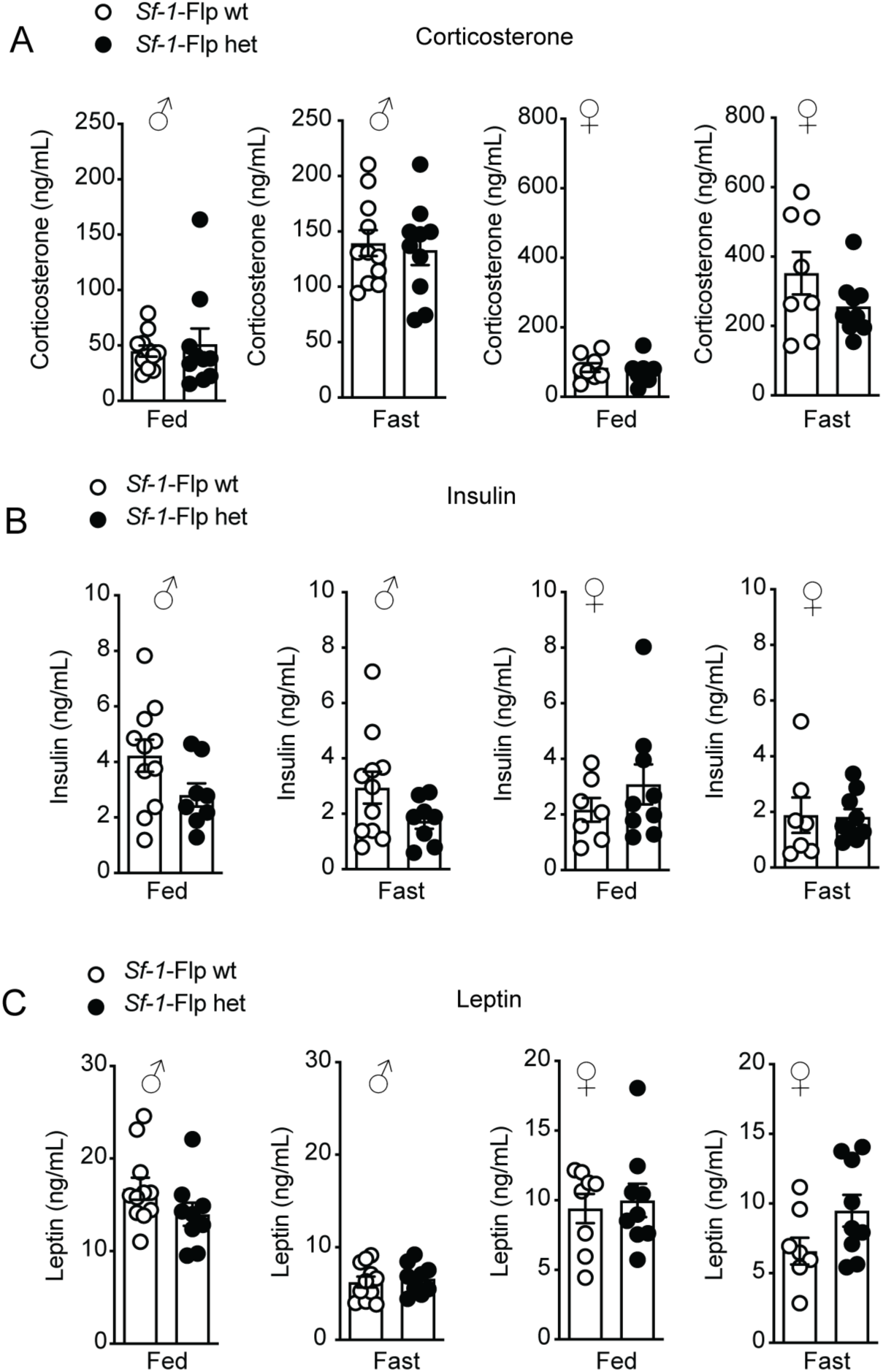
S*f-1*-Flp mice have normal responses of corticosterone, leptin, and insulin to fasting challenge. Blood (**A**) corticosterone, (**B**) insulin, (**C**) and leptin levels in fed and 24-hour fast conditions in *Sf-1*-Flp heterozygous male and female mice. Values are mean ± S.E.M.

**Supplemental Figure 4.**
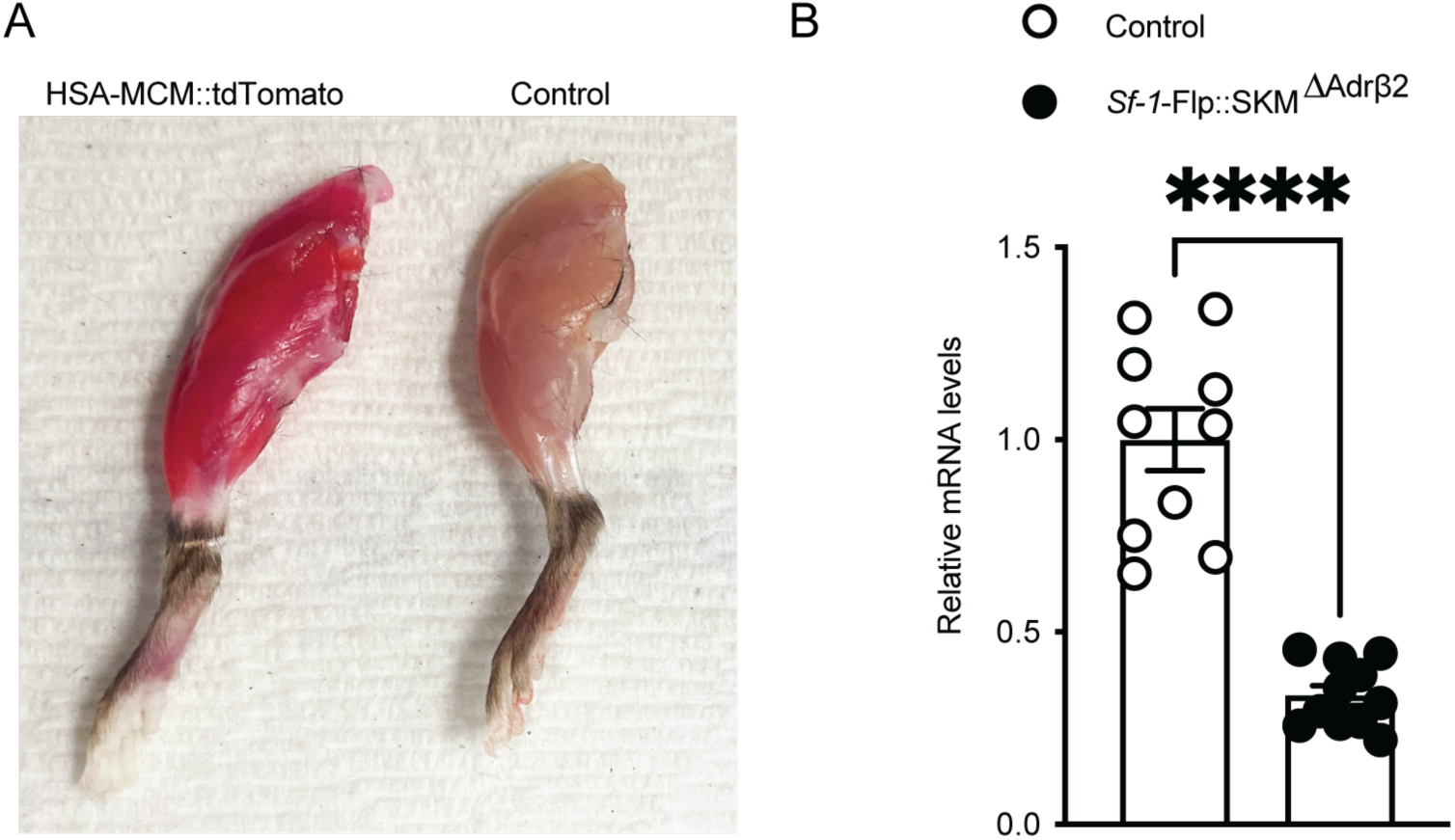
S*f-1*-Flp::SKM^ΔAdrβ2^ mice has lower Adrb2 mRNA expression in skeletal muscle. (**A**) Representative figure of lower limbs of HAC-MCM::Ai14^TB/-^ mice after 4 weeks tamoxifen (TAM) diet. A control group is composed of Ai14 mice. All mice were fed with 4 weeks TAM diet. (**B**) mRNA levels of *Adrb2* in TA skeletal muscle in *Sf-1*-Flp::SKM^ΔAdrβ2^ mice after 4 weeks tamoxifen diet. A control group is composed of *Sf-1*-Flp^+/-^::*Adrβ2*^flox/flox^ mice. All mice were fed with 4 weeks TAM diet. Values are mean ± S.E.M. **** p < 0.0001

